# Interferometric nanoparticle tracking analysis enables label-free discrimination of extracellular vesicles from large lipoproteins

**DOI:** 10.1101/2022.11.11.515605

**Authors:** Anna D. Kashkanova, Martin Blessing, Marie Reischke, Andreas S. Baur, Vahid Sandoghdar, Jan Van Deun

## Abstract

Extracellular vesicles (EVs) are increasingly gaining interest as biomarkers and therapeutics. Accurate sizing and quantification of EVs remain problematic, given their nanometer size range and small scattering cross-sections. This is compounded by the fact that common EV isolation methods result in co-isolation of particles with comparable features. Especially in blood plasma, similarly-sized lipoproteins outnumber EVs to a great extent. Recently, interferometric nanoparticle tracking analysis (iNTA) was introduced as a particle analysis method that enables determining the size and refractive index of nanoparticles with high sensitivity and precision. In this work, we apply iNTA to differentiate between EVs and lipoproteins, and compare its performance to conventional nanoparticle tracking analysis (NTA). We show that iNTA can accurately quantify EVs in artificial EV-lipoprotein mixtures and in plasma-derived EV samples of varying complexity. Conventional NTA could not report on EV numbers, as it was not able to distinguish between EVs and lipoproteins. iNTA has the potential to become a new standard for label-free EV characterization in suspension.

## Introduction

Pathological conditions can lead to a dysregulation of the circulating extracellular vesicle (EV) population, including in number and size. While this makes EVs an interesting target for biomarker studies on liquid biopsies, accurate sizing and enumeration of purified EV samples is typically hampered by the presence of co-isolated particles with similar properties as EVs. Lipoproteins are a main source of contamination when considering blood plasma. Intermediate-, very-low- and ultra-low-density lipoproteins (LPs) are of particular concern, as they outnumber EVs by several orders of magnitude and have an overlapping size range (30-1000 nm, therefore these will be designated in this paper as ‘large LPs’).^1^ Furthermore, their concentration in plasma varies with prandial status. Standard EV isolation methods such as differential ultracentrifugation or size-exclusion chromatography fail to remove a large percentage of large LPs.^2–4^

Limitations in commonly used, label-free EV quantitation systems, such as nanoparticle tracking analysis (NTA), do not allow a distinction between EVs and co-isolated large LPs, resulting in inaccurate sizing and overestimation of EV numbers.^4^ Differential labeling of both populations is hindered by the fact that most pan-EV labelling methods rely on general lipophilic or protein-binding dyes, which will also label LPs.^5^ Measuring a separate lysis buffer control has been suggested to verify particle origin, e.g. when analyzing EVs via flow cytometry, under the assumption that large LP particles would not be affected by this treatment.^6^ Recently, however, this approach was shown to be inadequate.^5^

Due to their different biochemical composition, EVs and large LPs differ in refractive index (RI). The RI is typically below 1.4 for EVs and above 1.4 for LPs.^7^ This results in distinct light-scattering properties, which can be used to separately analyze the two populations. However, until now this has only been successfully implemented for EVs larger than 200 nm, using a conventional flow cytometry-based approach.^7^ In another recent development, the different mechanical properties of EVs and lipoproteins have been used to discriminate them using atomic force microscopy.^8^ This technique however has low throughput and the substrate can affect the particle properties. A sensitive method that allows label-free discrimination of EVs and large LPs in solution would be highly beneficial.

We have recently developed a novel particle detection technology called interferometric NTA (iNTA).^9^ The defining feature of iNTA is the use of interferometric detection of scattering (iSCAT), in which the light scattered by a particle is interfered with light reflected by a cover glass, leading to enhanced sensitivity for small particle sizes.^10, 11^ In addition, recording trajectories at high frame rates (5 kHz) and measuring the amount of scattered light allows for accurate estimation of both size and RI for each individual particle. Previously, we verified the sensitivity and reproducibility of this method for EV detection, and found that it was able to distinguish EVs from urinary protein complexes of similar size.^9^ Here, we evaluated whether iNTA can be used to differentiate EVs from large LPs. Using mixtures of purified EVs and large LPs, as well as blood plasma samples purified by different methods, we show that iNTA can accurately measure the presence of bona fide EVs in plasma-derived samples of varying complexity.

## Methods

### Lipoprotein stock solutions

Low-density and very-low-density lipoproteins were acquired from Sigma (437644 and 437647). Ultra-low-density lipoproteins were purchased from Lee Biosolutions (194-14).

### Cell culture and generation of conditioned medium

Melanoma cell line SKMEL37 was cultured in RPMI 1640 (Life Technologies) supplemented with 10% fetal bovine serum (FBS), 1% penicillin–streptomycin (Lonza) and 2 mM L-glutamine (Lonza). Cells were washed 3 times with serum-free medium and cultured in RPMI supplemented with 1% EV-depleted FBS for 24 hours. FBS was depleted from EVs by ultracentrifugation (18 hours, 100000g). Conditioned medium (CM) (150 mL from ≈2E08 cells) was collected and centrifuged at 300g (10 minutes) and 2000g (20 minutes, 4°C). Next, CM was concentrated ≈300 times to 0.5 mL using a Centricon Plus-70 centrifugal filter device with a 10K nominal molecular weight limit (Merck Millipore).

### Blood collection

Venous blood was collected using a 21G needle in EDTA Vacutainers (BD367525, BD Biosciences). Pre-prandial blood of the healthy volunteer (male, 35 years old) was taken 12 hours after the last meal. Post-prandial blood was collected 3 hours after a meal with a caloric intake of ca. 1250 kCal (30% from fat). The melanoma patient included in this study (stage IB, female, 51 years old) was not in a fasting state at the time of blood collection. Collection of blood samples was approved by the Ethical Committee of the FAU Erlangen-Nürnberg (registration number 496_20B) and was performed in accordance with relevant guidelines. Donors have given written informed consent.

### Plasma generation

Platelet-free plasma was prepared by two serial centrifugations at 2500g for 15 minutes, first at room temperature and then at 4°C. The supernatant was aliquoted and stored at −80°C. All samples were processed within 2 hours after blood collection.

### Size-exclusion and dual-mode chromatography

SEC and DMC columns were prepared as described previously.^4^ Sepharose CL-4B (GE Healthcare) and Fractogel EMD SO3- (M) (Merck Millipore) resins were washed three times with PBS buffer. A nylon net with 20 μm pore size (NY2002500, Merck Millipore) was placed on the bottom of a 10 mL syringe (BD307736, BD Biosciences). For the SEC column, this was followed by stacking of 10 mL washed Sepharose. For the DMC column, 2 mL of Fractogel was stacked first, followed by careful layering of 10 mL Sepharose on top. After adding 0.5 mL plasma sample, the column was eluted by constant addition of fresh PBS. After discarding the void volume, which was 3 mL for SEC and 3.5 mL for DMC, 2 mL of EV-containing eluate was collected and concentrated to 100 μL using Amicon Ultra-2 10K filters (Merck Millipore).

### Density gradient

A discontinuous OptiPrep density gradient (DG) was constructed as described previously,^12^ with some modifications. Solutions of 5, 10, 20 and 40% iodixanol were made by mixing appropriate amounts of homogenization buffer (0.25M sucrose, 1mM EDTA, 10 mM Tris-HCL, pH 7.4) and OptiPrep stock solution (60% (w/v) aqueous iodixanol, Axis-Shield, Oslo, Norway). The gradient was formed by layering 3 mL of 40%, 3 mL of 20%, 3 mL of 10% and 2.5 mL of 5% solutions on top of each other in a 12 mL open top polyclear tube (Seton Scientific). 500 μL of sample was overlaid onto the top of the gradient, which was then centrifuged for 18 hours at 100000g and 4°C (TH641 rotor, Thermo Fisher). Afterwards, gradient fractions of 1 mL were collected from the top of the gradient and their density was estimated using a refractometer (Abbemat 200, Anton Paar). Fractions 7 and 8, corresponding to an EV density of 1.1-1.2 g/mL, were pooled and used for subsequent SEC-based separation of EVs from the iodixanol polymer, as described above and previously.^13^ EV-containing fractions (F4-7) were pooled, concentrated to 100 μL, aliquoted into Protein LoBind tubes (Eppendorf), and stored at 4°C (maximum 48 hours, for comparative NTA and iNTA measurements) or −80°C. For DG-based EV isolation of melanoma patient plasma, the gradient procedure described above was preceded by a SEC separation, based on Vergauwen et al.^14^ A total of 6 mL of plasma sample was divided over 3 SEC columns (2 mL of plasma per SEC column) followed by elution by PBS. Individual fractions of 1 mL were collected. Fraction 4-6 of three different columns were pooled and concentrated to 0.5 mL using an Amicon Ultra-15 10K filter (Merck Millipore). This was then placed on top of the gradient.

### Nanoparticle tracking analysis (NTA)

A Zetaview PMX-110 instrument, equipped with a 405 nm laser, was used to measure particle concentrations. Before sample measurement, the instrument was calibrated using 100 nm polystyrene beads diluted in water according to manufacturer’s instructions. Cell temperature was maintained at 25°C for all measurements. Samples were diluted to an appropriate concentration in PBS, in a total volume of 1 mL. Eleven cell positions were scanned for each measurement cycle, with video recorded at 30 frames per second. Additional capture settings were: gain 719.52, shutter 50, minimum trace length 15. For measuring 40 nm and 60 nm PS beads, the gain was increased to 888.12. ZetaView software version 8.05.12 was used to analyze the recorded videos with the following settings: minimum brightness 25, maximum brightness 255, minimum area 5, and maximum area 200.

### Interferometric nanoparticle tracking analysis (iNTA)

An iNTA measurement setup^9^ was employed for the measurements. Measurements were done both in chambered coverglasses (IBIDI μ-Slide 18 well) and in home-made PVC chambers.^15^ The PVC chamber’s bottom coverglasses were passivated with mPEG2000-Silane (Laysan Bio). For that purpose, borosilicate coverglasses (VWR) were sonicated in 2% Hellmanex III (Hellmanex Analytics) and milliQ (milliPore) subsequently for 10 minutes each. Clean coverglasses were dipped into ethanol and isopropanol before blow drying with nitrogen gas. The coverglasses were plasma cleaned (10 minutes in oxygen plasma at 500 W) and then incubated in 10 mg/mL mPEG2000-Silane at 50 °C. mPEG200-Silane was dissolved in PEG buffer (95% Ethanol (v/v), 5% milliQ, pH was set to 2.0 with 1M HCl). When the buffer fully evaporated the coverglasses were sonicated for another 10 minutes in milliQ and blow dried with nitrogen gas. Passivated coverglasses were used the same day. The procedure for passivating the chambered coverglasses is similar, however the milliQ, ethanol and isopropanol cleaning steps were skipped.

For the measurement the microscope focus was set at 1 μm above the coverglass. Videos of particles diffusing in 0.5-1.0 μl droplets of fluid on the coverglass (Supp. Fig. 1) or in 50-100 μL volumes of fluid inside individual IBIDI wells were recorded for 10 minutes using pylablib cam-control (https://github.com/SandoghdarLab/pyLabLib-cam-control). They were analyzed by applying median background correction and radial variance transform.^16^ Particles were tracked using trackpy python package.^17^ The particle size was extracted from its diffusion constant. The particle’s scattering cross-section was extracted from its iSCAT contrast using previously performed calibration.^9^ The effective RI was then calculated from the size and scattering cross-section.

To enable a direct comparison of particle concentrations measured by NTA and iNTA, we performed a calibration measurement using a dilution series of 60 nm polystyrene beads (Supp. Figure 2). This allowed us to extract a calibration factor for iNTA, which we used for converting a number of particle trajectories detected in a given time to a concentration in terms of number of particles per millilitre.

### ELISA

Human Apolipoprotein B Quantikine ELISA Kit (R&D Systems, DAPB00) was used according to manufacturer’s instructions. Standards and samples were assayed in duplicate. CD63 ELISA (Abcam, ab275099) was performed according to manufacturer’s instructions. Due to limitations in available sample amount, samples were only assayed once.

### Western blotting

Equal volumes of EV samples were suspended in non-reducing sample buffer (0.05 M Tris-HCl (pH 6.8), 10% glycerol, 2% SDS, 1% bromophenol blue) and boiled for 5 minutes at 95°C. Proteins were separated by SDS-PAGE (SDS polyacrylamide gel electrophoresis), transferred to nitrocellulose membranes, blocked in 5% nonfat milk in PBS with 0.5% Tween-20, and immunostained overnight at 4°C using the following primary antibodies in a 1/1000 dilution in TBST: CD9 (BD555370), CD63 (BD556019), CD81 (BD555675). Blots were developed using the SuperSignal West Femto reagent (Thermo Fisher) and visualized on an Amer-sham 600 system (GE Healthcare). CD9 and CD81 were stained on the same membrane, which was stripped for 10 minutes using Restore Western blot stripping buffer (Life Technologies) in between staining.

### EV-TRACK knowledgebase

We have submitted all relevant data of our experiments to the EV-TRACK knowledgebase (EV-TRACK ID: EV220321).^18^

## Results

### iNTA allows simultaneous size and refractive index estimation of EVs and large LPs

First, to compare the performance of NTA and iNTA for analyzing biological nanoparticles, we measured the concentration and size of density gradient-purified cell culture EVs and commercial LP solutions. Given the interest in using EVs for cancer detection, the SKMEL37 melanoma cell line was selected as EV source. Measured particle concentrations for EVs and VLDLs were higher in iNTA compared to NTA, while for ULDLs they were similar (Table 1). LDLs, a class of LPs which are typically smaller than EVs (<30 nm^19^), were not detected accurately by NTA nor iNTA (Supp. Figure 3). Size distributions as measured by iNTA were narrower and shifted towards a smaller size, compared to NTA (Table 1 and Fig. 1a). Indeed, using polystyrene spheres and silica beads of different sizes, we found that NTA generally returns a broader size distribution with an additional offset for small/weakly scattering particles (Supp. Fig. 4), as was also shown previously by others.^20–22^ Taken together, these data show that iNTA has an increased sensitivity for smaller particles, whether artificial or biological, consistent with previous data.^9^

**Figure 1:**
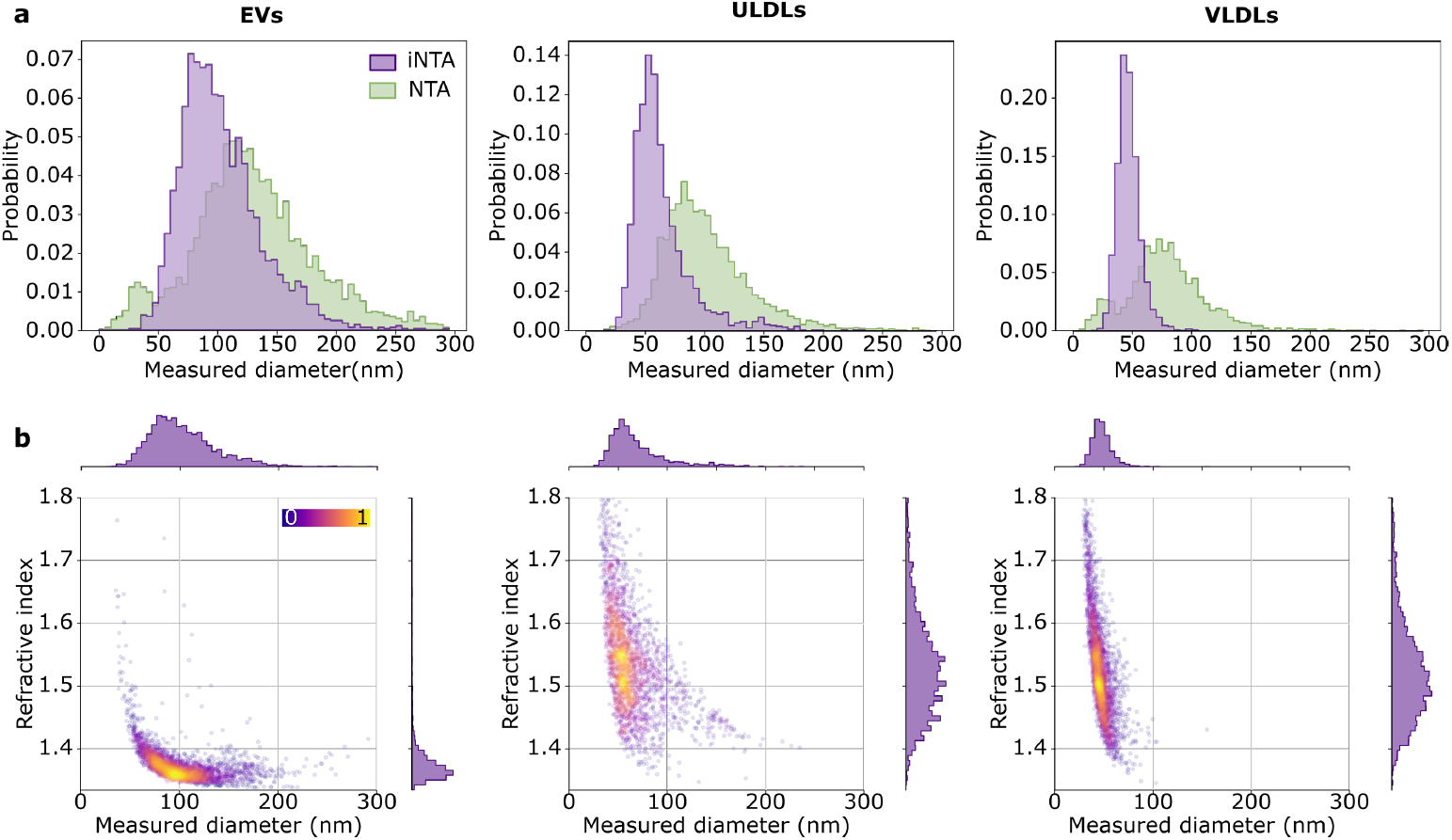
iNTA is more sensitive than NTA for smaller-sized EVs and LPs, and enables refractive index measurements. **(a)** EV, ULDL and VLDL size distributions as measured by NTA (green) and iNTA (purple). **(b)** Size-RI plots for EV, ULDL and VLDL measured by iNTA. Color bar indicates the local density of points.

**Table 1:**
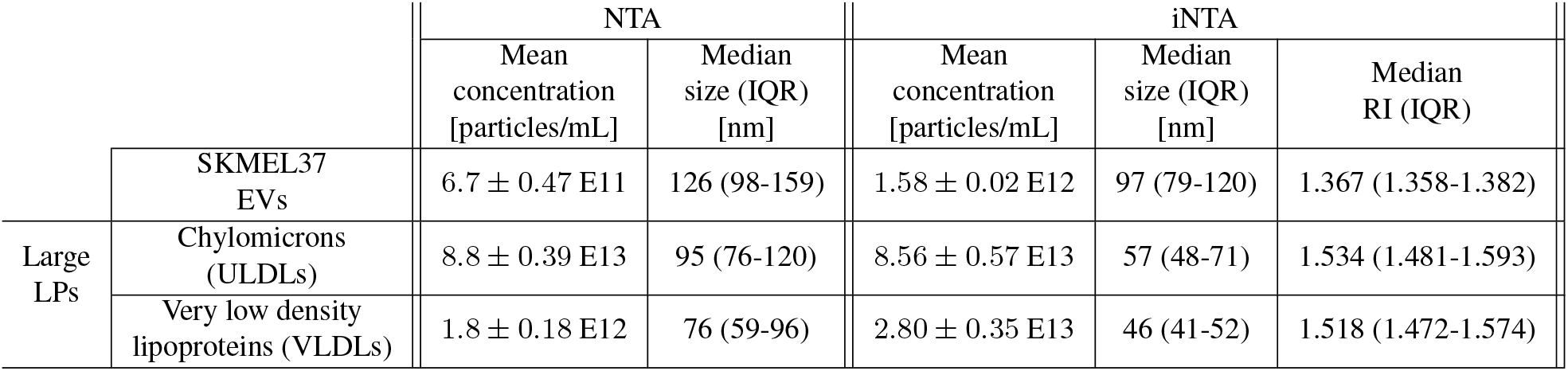
Mean concentration and median size including interquartile range (IQR) of different classes of nanoparticles measured using NTA and iNTA. Refractive index is also shown for iNTA. For concentration, the standard deviation of three successive measurements is indicated.

Next, we aimed to differentiate EVs and large LPs based on their distinct light scattering properties due to a difference in refractive index (RI).^7^ We found that the mean scattering intensity as recorded for each particle by NTA could not be used to distinguish EVs from large LPs (Supp. Fig. 5). For iNTA, however, the amount of scattered light for each particle is measured more precisely, using iSCAT contrast (Supp. Fig. 4).^9^ This allowed us to derive distinct RIs for EVs (median RI of 1.367) and large LPs (median RIs of 1.534 for ULDLs and 1.518 for VLDLs) (Table 1). These values agree with those reported in literature.^23, 24^ Aside from a small overlapping region of RI ≃ 1.4 and size 50-100 nm, EVs and large LPs appear as separate populations in size-RI plots generated by iNTA (Fig. 1b). Based on these measurements, a machine learning classifier was constructed to discriminate EVs and LPs (Supp. Fig. 6).^25^ For each particle with a given size, scattering cross-section and RI, the classifier returns the probability of it being an EV or a large LP. Using an 80% confidence threshold, iNTA measurements enable us to classify more than 85% of measured particles as either EVs or large LPs. This threshold was thus selected to be used in our further analyses.

### Mixed populations of EVs and LPs are discriminated by iNTA

Given the ability of iNTA to simultaneously determine RI and size for both EVs and LPs, we tested whether this would also allow accurate estimation of EV numbers in mixtures with LPs. We started with a mixture of SKMEL37 EVs with VLDLs in 3:1, 1:1 and 1:3 volume ratio. Roughly equal numbers of EVs and VLDLs as estimated by iNTA were chosen as starting material.

With iNTA, we measured equal total particle numbers for all mixtures, as expected (Fig. 2a). The median size of the mixtures gradually decreased with increasing proportion of VLDLs, which were indeed measured to be almost two times smaller than EVs. NTA detected more than twofold fewer EVs and tenfold fewer VLDLs than iNTA (Fig. 2b). As a result, a stepwise decrease in total particle concentration from EVs-only to VLDLs-only could be observed. However, even though VLDLs were measured to be around 1.5 times smaller than EVs, the median size remained almost constant in all mixtures containing EVs. This confirms that NTA is biased towards larger particles which scatter more light,^20^ contributing to the fact that it could not distinguish between EVs and VLDLs in these mixtures (Supp. Fig. 7). Even using iNTA, the broad size distributions of VLDLs and especially EVs did not allow for quantitative discrimination of the two particle types based on size alone. However, by simultaneously determining the RI for each particle, distinct populations could be resolved on size-RI plots using our machine learning classifier (Fig. 2c, EVs indicated in red and LPs in blue). Indeed, this allowed us to reproduce the different ratios of EVs and VLDLs (Fig. 2d - left panel). We were also able to extract size and RI of the individual EV and LP populations in the mixtures (Fig. 2d - right panels). Technical replicates of these measurements showed only minor differences, demonstrating the robustness of iNTA for simultaneous size and RI determination of individual biological nanoparticles (Supp. Fig. 8). To show that iNTA can also decipher more complex mixtures, we spiked four different quantities of EVs – spanning concentrations of 6% up to 54% – into a solution containing ULDLs, VLDLs and LDLs. Also in these samples, iNTA was able to accurately discriminate EVs from LPs (Fig. 2e and Supp. Fig. 9). The same mixtures measured by NTA again showed a good correspondence between observed and expected total particle concentrations, but no discrimination of the different particle types (Supp. Fig. 10).

**Figure 2:**
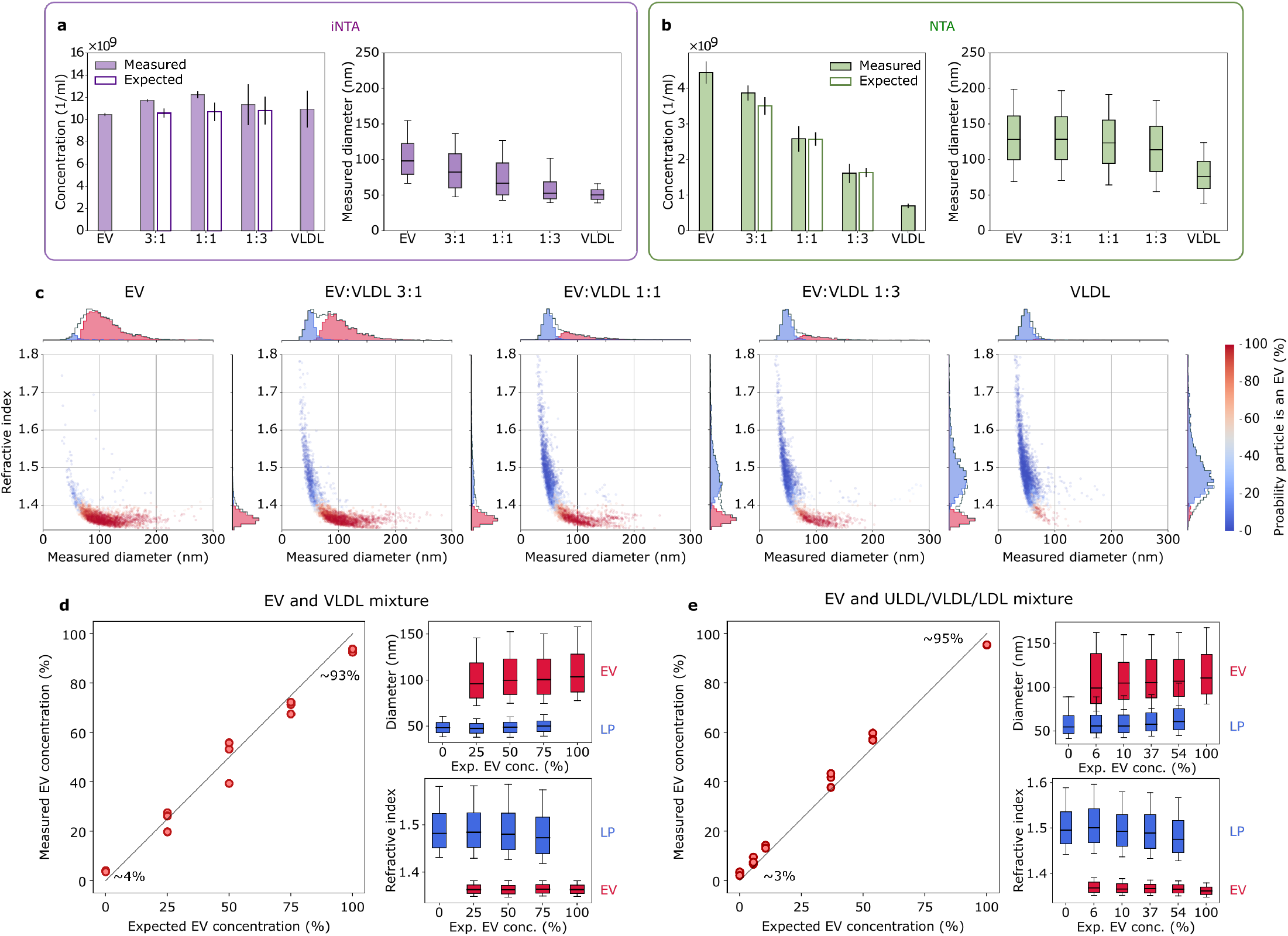
iNTA but not NTA can discriminate EVs and LPs in artificial mixtures. **(a)** iNTA and **(b)** NTA analysis of different EV-VLDL mixtures showing mean particle concentration (filled bars indicate the measured concentration while unfilled bars indicate the expected concentration based on measurements of pure EV and VLDL samples, standard deviation is shown) and median size (box boundaries are 25-75 percentile, horizontal line is median, whiskers indicate 10-90 percentile) (n=3). Concentration for iNTA measurements is calculated by multiplying the number of trajectories measured in 10 minutes by a calibration factor determined previously (see Methods). **(c)** Representative size-RI plots for the different EV-VLDL mixtures, labeled using a random forest classifier. Color indicates probability for the particle to be an EV. Only particles with greater than 80% probability of being an EV or less than 20% probability of being an LP are included in the 1D histograms. **(d)** Left: Measured relative EV concentration plotted vs. expected relative EV concentration. The measured concentration is calculated as *C*_EV_ / (*C*_EV_+*C*_VLDL_), where *C* denotes particle concentration determined from plots in (c) using the random forest classifier. The numbers indicate the measured relative EV concentration when no EVs were expected (~ 4%) and when only EVs were expected (~ 93%). Right: The extracted diameter and refractive index of EVs (red) and lipoproteins (blue). **(e)** Same as (d), but using a mixture of EVs with ULDLs, VLDLs and LDLs.

Summarizing the iNTA data, we found that 3-4% of particles were misclassified as EVs in pure LP samples, while 5-7% of particles were misclassified as LPs in pure EV samples. This demonstrates that iNTA, unlike NTA, is able to accurately detect bona fide EVs in complex mixtures with LPs.

### iNTA is not affected by residual lipoprotein contamination in plasma-derived EV samples

In order to evaluate the performance of iNTA to measure EVs in real-world samples, including the effect of variable plasma lipid composition, we collected blood plasma from a healthy volunteer in preprandial or postprandial state (i.e. before versus 3 hours after consuming a fatty meal). ELISA measurements confirmed an increase in the plasma concentration of ApoB, a marker for ULDLs, VLDLs and LDLs, after meal consumption (Supp. Fig. 11). We also compared two EV enrichment methods, each known to remove LPs to a different extent. We used either size-exclusion chromatography (SEC), where all LPs with a similar size as EVs will be collected in the same fraction, or dual-mode chromatography (DMC), a method capable of removing >99% of ApoB-containing LPs.^4^

As expected, ELISA measurements showed an increased ApoB concentration in post-versus preprandial EV samples, as well as in those obtained by SEC compared to DMC (Fig. 3a). iNTA particle counts showed a similar result, increasing after meal consumption and SEC samples showing higher concentrations compared to DMC (Fig. 3b). Similar trends in total particle concentrations were observed for conventional NTA (Supp. Fig. 12). Extracting the nature of the particles from the size-RI plots generated by iNTA, we found that this increase in particle concentration is exclusively due to large LPs (Fig. 3c). Indeed, the relative EV concentrations were below 4% for all unspiked samples (Fig. 3d), consistent with the misclassification rate of LPs that we determined previously. The low level of EVs in samples from this particular individual was confirmed using a CD63 ELISA, with all measurements being below limit of detection (Supp. Fig. 13).

**Figure 3:**
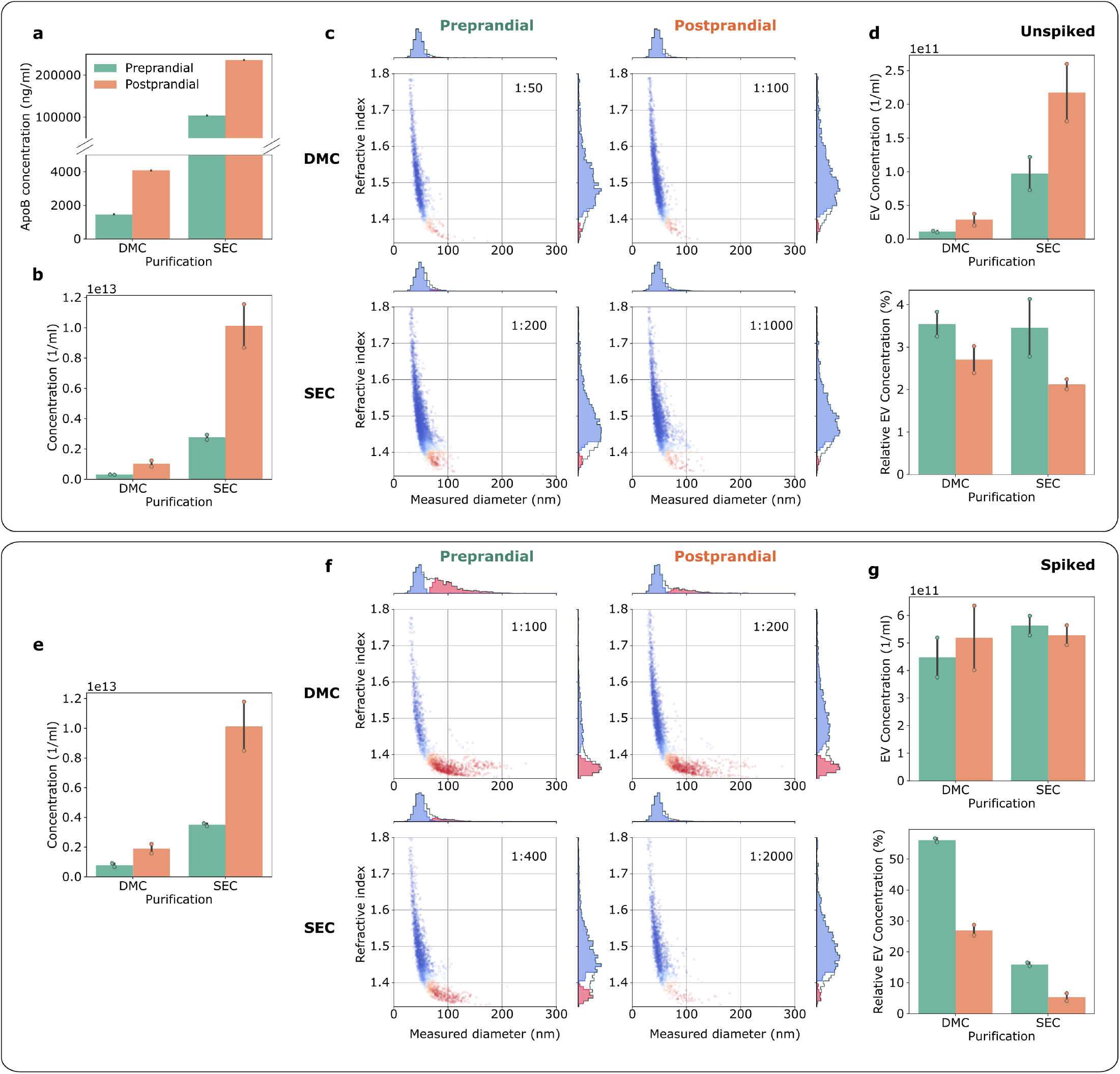
iNTA can accurately measure EVs in plasma-derived samples with variable LP background. **(a)** ApoB ELISA measurements in DMC- and SEC-processed samples (left and right), pre- and post-prandial (green and orange). **(b)** Total particle concentrations measured by iNTA. **(c)** Representative size-RI plots after machine learning processing (particles classified as EVs are indicated in red and LPs in blue). The numbers in the upper right corner indicate sample dilution factor by PBS prior to iNTA measurements. **(d)** EV concentration (top) and relative EV concentration (bottom) as determined by iNTA. **(e)**, **(f)** and **(g)** Same as (b), (c) and (d) but for samples spiked with 5E11/mL SKMEL37 EVs. Mean and standard deviation are shown for all bar graphs (n=2).

To show that iNTA is able to accurately detect EVs in these samples, we decided to make use of an exogenous EV spike. SEC and DMC samples were spiked with 5E11 particles/mL cell culture EVs and analyzed by iNTA, as shown in Fig. 3e,f. Despite the differences in LP background, iNTA was capable of extracting the spiked EV concentration from all samples with only minimal error (Fig. 3g). The same samples measured by NTA showed an increase in total particle concentration, however the spiked EV concentration could not be estimated (Supp. Fig. 12). This demonstrates that EVs can be quantified by iNTA in plasma-derived samples of varying LP complexity, regardless of prandial status and resulting changes in LP level.

Finally, to show that iNTA is able to measure endogenous EVs derived from a lipid-rich sample, we collected plasma from a non-fasting melanoma patient. EV concentration was expected to be increased compared to healthy plasma, as shown previously for melanoma.^26^ EVs were isolated using SEC and DMC, as before, and also by a density gradient-based protocol (DG). This protocol is known to result in EVs of high purity.^14^ A 12-times higher plasma sample input was used for the DG procedure as for SEC/DMC (6 mL versus 0.5 mL). We could confirm the presence of EVs in all three samples using Western blotting for CD9, CD63 and CD81 (Fig. 4a and Supp. Fig. 14). The strongest signals were present in DG, as expected due to the higher sample input. To estimate the presence of LPs, we also measured ApoB concentration in these samples (Supp. Figure 11). The plasma sample was found to contain around 30%more ApoB than the postprandial healthy volunteer, indicating very high LP levels. SEC, DMC and DG each removed ApoB to a different extent, by around 25%, 99% and 100%, respectively. These same samples were then measured by iNTA. The SEC-processed sample showed a EV concentration of 8.8E11 particles/mL, however with a relative concentration of only 3.7% due to high LP content, *i.e*. within the range of classification error (Fig. 4b). DMC-based EV enrichment was not sufficient to avoid large LPs outnumbering EVs. Nevertheless, iNTA could still identify a clear EV population with a concentration of 9.8E10 particles/mL and relative concentration of 7.2% (Fig. 4c). For DG, iNTA detected a clear EV population with no background of large LPs (Fig. 4d, EV concentration of 9.0E11 particles/mL and relative concentration of 97.7%). Using these data and taking into account a previously-published recovery rate of 25% for the DG protocol,^27^ the original EV concentration in this plasma sample was estimated to be around 6E10 particles/mL. The same calculation was made for the DMC and SEC measurements, using recovery rates of 33% and 80%, respectively.^4^ While the calculated value based on DMC agreed with that of ODG (5.9E10 particles/mL), SEC was off by a factor of four (2.2E11 particles/mL) due to the overwhelming presence of large LPs in this sample. This confirms that iNTA is able to accurately quantify endogenous EVs in large LP-containing samples up until an EV to LP ratio of around 1 to

**Figure 4:**
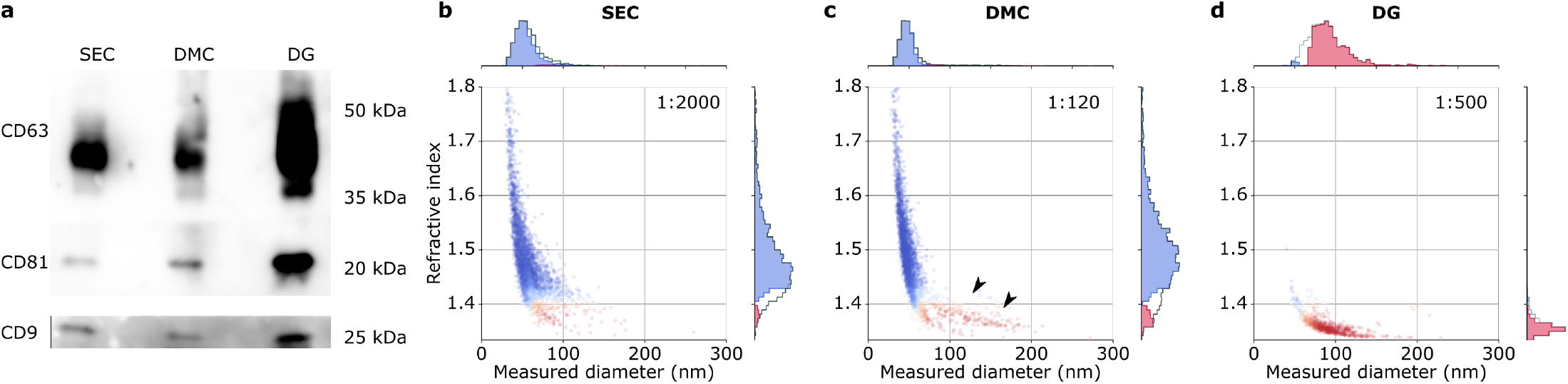
iNTA of EV samples enriched from lipemic melanoma patient plasma by SEC, DMC or DG. **(a)** Western blot of EV-associated tetraspanins. Equal volumes of EV sample were loaded, with DG (6 mL) having a 12x higher plasma sample input than SEC and DMC (0.5 mL). Size-RI plots of EVs enriched by **(b)** SEC; **(c)** DMC, arrowheads indicate the EV population; **(d)** DG.

## Discussion

We have demonstrated that iNTA can discriminate EVs from large LPs in a label-free manner, through simultaneous determination of particle size and refractive index. We have constructed a machine learning classifier for EVs and large LPs, and experimentally validated that EV concentration can be quantified accurately, also when outnumbered by large LPs by up to 15x. iNTA measurements are highly sensitive, fast (10 minutes), and require only a small sample volume (≤1 μl). In addition, our work highlights the limited applicability of conventional NTA for determining EV concentration in complex mixtures of biological particles, such as blood plasma EV samples obtained by SEC. While total particle concentrations of EV-LP mixtures corresponded well to theoretical values, the relative contributions of EVs and LPs could not be determined by NTA. As large LPs outnumber EVs in many cases, NTA data have to be interpreted with care. Furthermore, NTA does not reach a sufficient sensitivity towards smaller EVs (or LPs) in a heterogeneous population, whereas iNTA generally reaches a smaller size range.

Despite the clear advantages of iNTA, a few challenges remain. The reliance on RI to distinguish between EVs and LPs is inherently limited by their RI overlap. Especially for LPs, we found a broad RI distribution. While LPs are indeed reported to be highly heterogeneous,^28^ this is also due to technical factors. First, for smaller particles, the trajectories are shorter and therefore the error on particle size increases. Second, for particles with a small scattering cross-section, the relative error on iSCAT contrast is increased. Since RI is calculated from a combination of size and iSCAT contrast, the errors in both lead to a spread in RI. This also causes the narrowing of the RI distribution with increasing particle size as seen in Supp. Fig. 4. As a result of the overlap, about 6% of EVs are misclassified when using an 80% confidence threshold for our machine learning classifier. The accuracy of iNTA for absolute EV quantification will therefore vary with the relative abundance of EVs and large LPs in a given sample. This limitation can be observed in the iNTA measurement of SEC EVs of the highly lipemic melanoma sample. While the number of particles classified as EVs was below the margin of error, Western blotting did identify similar EV marker presence as in the DMC sample.

Additional changes to iNTA could help to improve its performance further. In the current study, iNTA was implemented using static measurements inside nano/microliter droplets. Having a dynamic measurement, *e.g*. using a constant flow, would produce a more representative image of the sample. In addition, it could help to more accurately quantify lower numbers of EVs and cut down on measurement time. Another limitation concerns *absolute* concentration measurements. In this study, particle concentration measured by iNTA was calibrated using an NTA measurement of 60 nm PS beads. This strategy is suboptimal, given the known limitations of NTA for concentration measurements.^20, 22, 29^ It is thus likely that the measured concentrations, e.g. for the melanoma patient, still do not represent the true number. A detailed study of independently using iNTA for absolute particle concentration determination is underway and will be reported elsewhere. Further possible improvements to iNTA include increasing the measurement sensitivity by using higher laser power and shorter wavelengths, adding measurements of zeta-potential on a single particle level to further improve differentiation between particle types, and expanding its capabilities for detecting labeled EV subpopulations by adding fluorescence filters. A significant advantage compared to current single particle fluorescence approaches would be that mislabeled non-EV particles can be filtered out using RI measurements.

Overall, iNTA outperformed conventional NTA. Indeed, we envision it could have a broad impact on EV research as a new standard for label-free EV sizing and quantification. It could also facilitate the translational application of EVs, for example by improving EV-based biomarker normalization and enabling superior quality control of EV products such as reference materials and therapeutics.

## Acknowledgements

We thank the Max Planck Society and the Bundesministerium für Bildung und Forschung for financial support. A.K. was supported by a Christiane Nüsslein-Volhard-Stiftung Fellowship and Alexander von Humboldt-Stiftung Postdoctoral Fellowship.

## Data availability

The data that support the findings of this study are available from the corresponding authors upon request.

