## Supplemental Information for "Interferometric nanoparticle tracking analysis enables label-free discrimination of extracellular vesicles from large lipoproteins"

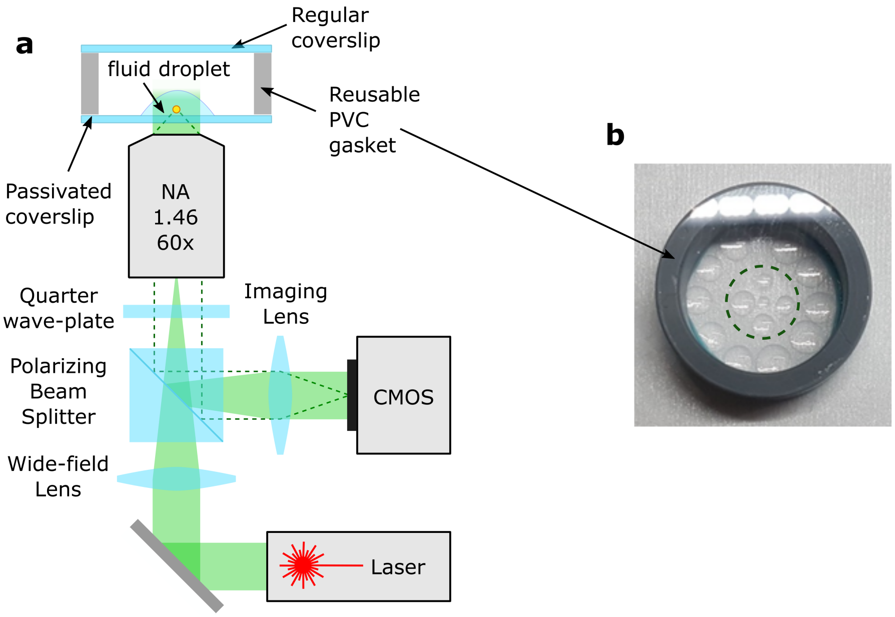


**Figure S1: iNTA measurement setup.** **(a)** Wide-field iSCAT setup for tracking freely diffusing particles. Linearly polarized light from a laser passes a polarizing beam splitter followed by a $\lambda/4$ plate that renders its polarisation circular. A wide-field lens ($f = 400$ mm) focuses the light at the back focal plane of the objective. An imaging lens ($f = 500$ mm) projects the reflected (solid green area) and scattered (dashed line) light on a CMOS camera chip.**(b)** A PVC chamber used for the measurements. A reusable PVC gasket is glued using dental glue to a coverslip passivated with Sil-PEG (see Methods). The droplets containing the sample are positioned in the center of the coverglass (inside dashed green circle), here shown with volumes ranging from 200 nl up to 2 $\mu$l. For our measurements, we worked with volumes from 500 nl up to 1 $\mu$l. On the perimeter several large droplets of PBS are positioned to serve as a "reservoir" to keep humidity inside the chamber high and prevent evaporation. Another coverslip is positioned on top of the PVC gasket.

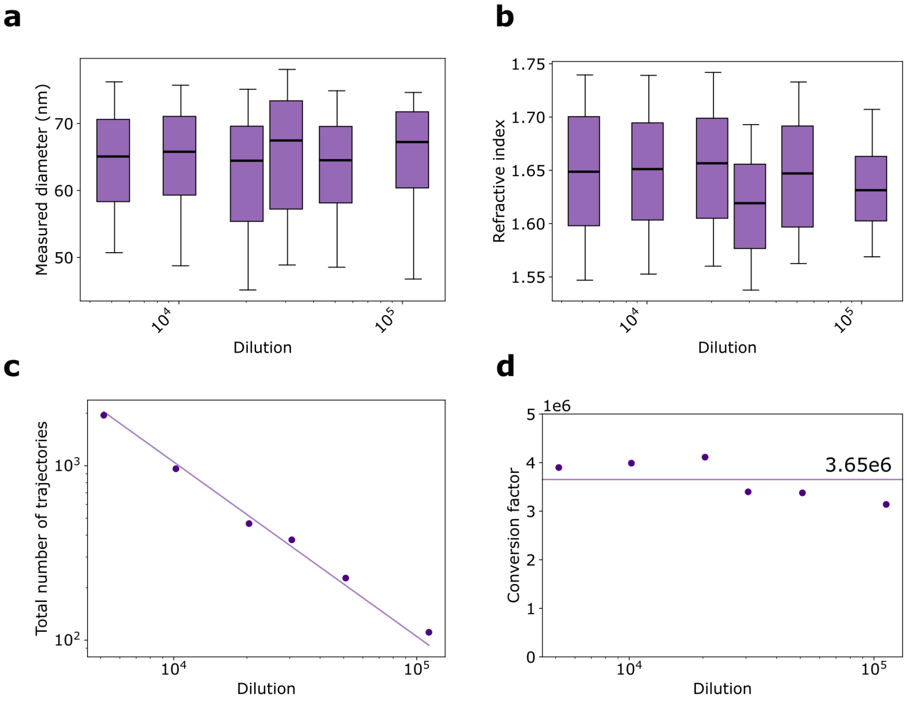


**Figure S2: iNTA particle concentration calibration.** A sample of 60 nm polystyrene spheres was measured with iNTA at different dilutions factors (DF) from 5000x to 100,000x. The measured **(a)** size and **(b)** refractive index of particles at different dilutions demonstrate the robustness of the method (median and 10-90 percentile (whiskers)). **(c)** the number of trajectories ($N_{\mathrm{traj}}$) recorded over 10 minutes for different dilutions. Only particles that cross the focal plane (es evidenced by the width of the point spread function) are included. **(d)** Using the NTA measurement of particle concentration ($C_{\mathrm{NTA}}$) for this sample we calculate the conversion factor as $C_{\mathrm{NTA}}/(N_{\mathrm{traj}} \times DF) = 3.65\mathrm{E}6$, which allows us to convert the number of trajectories into particle concentration.

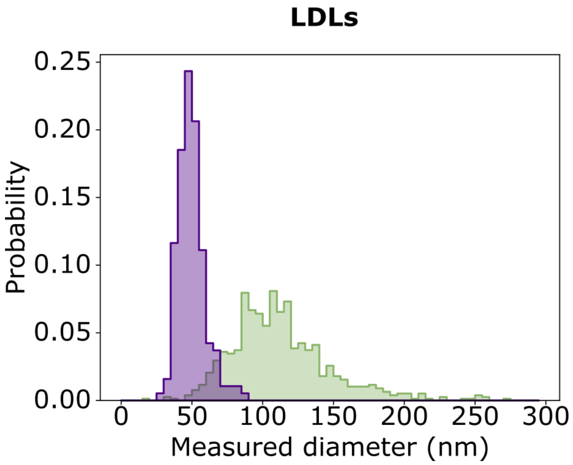


**Figure S3: Low density lipoprotein size distribution**. Measurement performed using NTA (green) and iNTA (purple). Both techniques measured particles of larger size than expected for LDLs (median size of 108 nm for NTA and 48 nm for iNTA), possibly representing large LPs contaminating the LDL preparation.

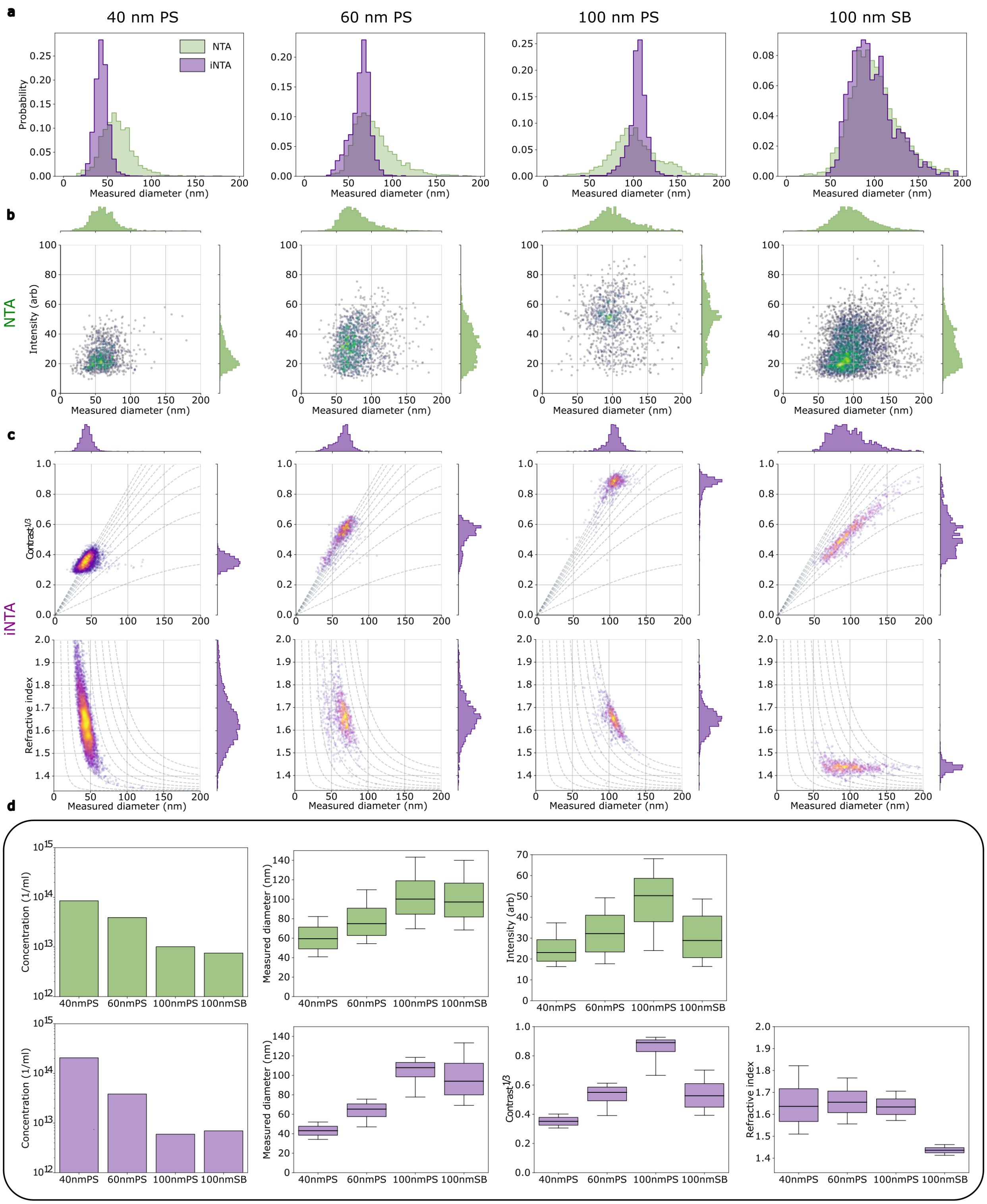


**Figure S4: Performance comparison of NTA and iNTA using synthetic beads.** **(a)** 40, 60, 100 nm polystyrene spheres (PS) and 100 nm silica beads (SB) size distributions as measured by NTA (green) and iNTA (purple) **(b)** NTA plots of intensity vs. size. Blue tones indicate lower density of points and yellow tones indicate higher density of points. **(c)** iNTA plots of contrast vs. size (top row) and refractive index vs. size (bottom row). Purple tones indicate lower density of points and yellow tones indicate higher density of points. In the top row dashed gray lines indicate the constant refractive index ranging from 1.34 to 1.66 in steps of 0.04. In the bottom row the dashed line indicate lines of constant contrast$^{1/3}$ from 0.1 to 0.9 in steps of 0.1. Note that the extracted diameter and contrast are correlated, despite being measured independently. This confirms that both are extracted correctly. The effective refractive index for small polystyrene spheres has a broader size distribution than for large polystyrene spheres and for silica beads. This is caused by the closer spacing of refractive index lines on size/contrast plot for smaller particles and larger RI. **(d)** Summary plots for NTA (top) and iNTA (bottom). Concentration is displayed as mean, diameter/contrast/intensity as median and 10-90 percentile (whiskers).

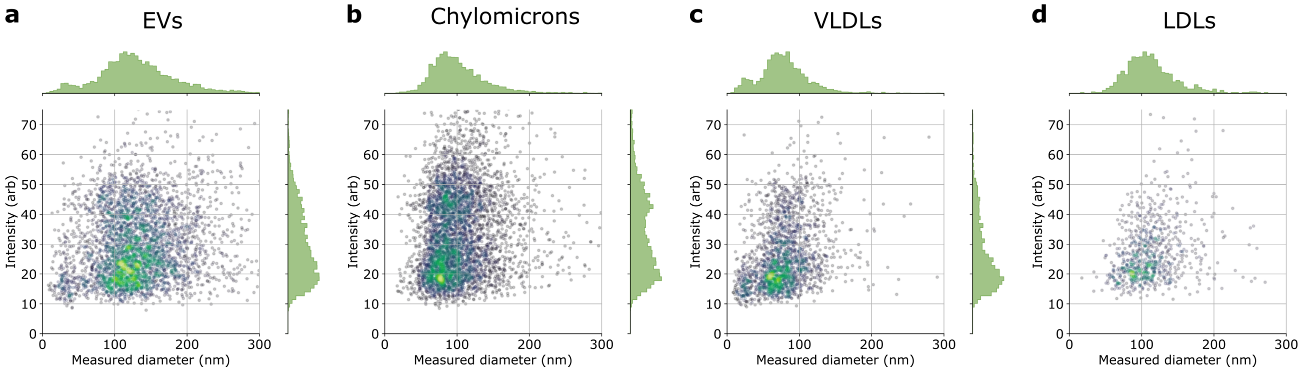


**Figure S5: NTA measurements of EVs and lipoproteins.** The scattered intensity returned by the instrument is plotted vs. the measured diameter. The blue tones correspond to lower point density while the yellow tones indicate higher point density. The samples are **(a)** EVs, **(b)** ULDLs, **(c)** VLDLs, and **(d)** LDLs.

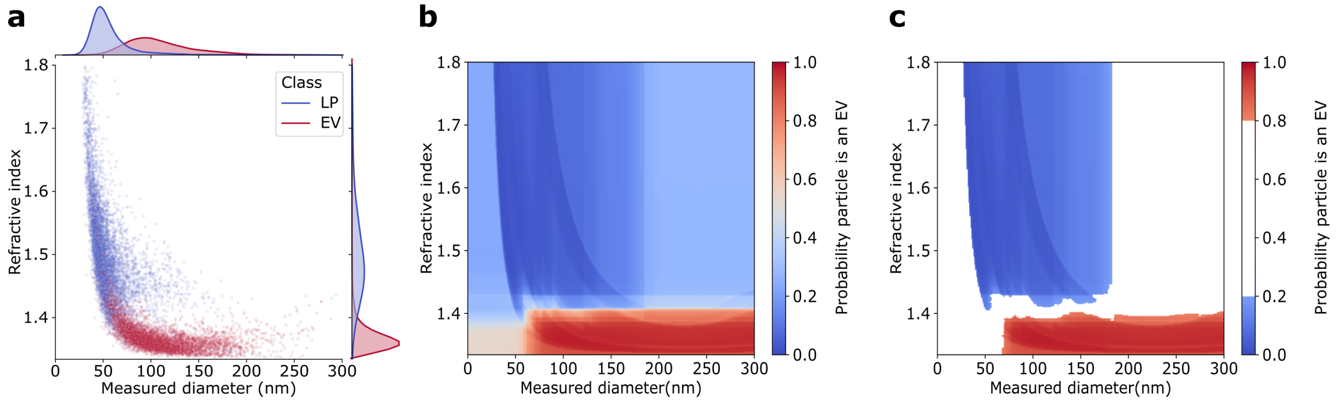


**Figure S6: Random forest particle classifier to differentiate EVs and lipoproteins.** **(a)** Training dataset with two separate measurements of EVs and lipoproteins. **(b)** Resulting probability map returned by the classifier trained using python scikit-learn package. The features used by the classifier are particle size, maximum iSCAT contrast to the power of 1/3 and refractive index. The parameters are as follows: 500 estimators, balanced class weight, 10 samples required to be at a leaf node minimum and maximum depth of the tree of 5. The other parameters are kept at their default values. **(c)** Same map as in (b) but only regions with $\geq 80\%$ confidence of particle belonging to either class are plotted.

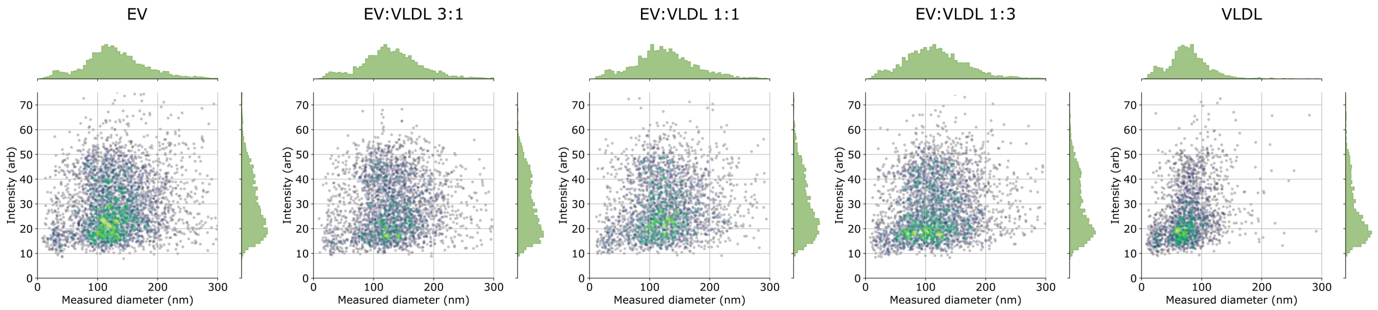


**Figure S7: NTA measurements of EV-VLDL mixtures.** The blue tones correspond to lower point density while the yellow tones indicate higher point density.

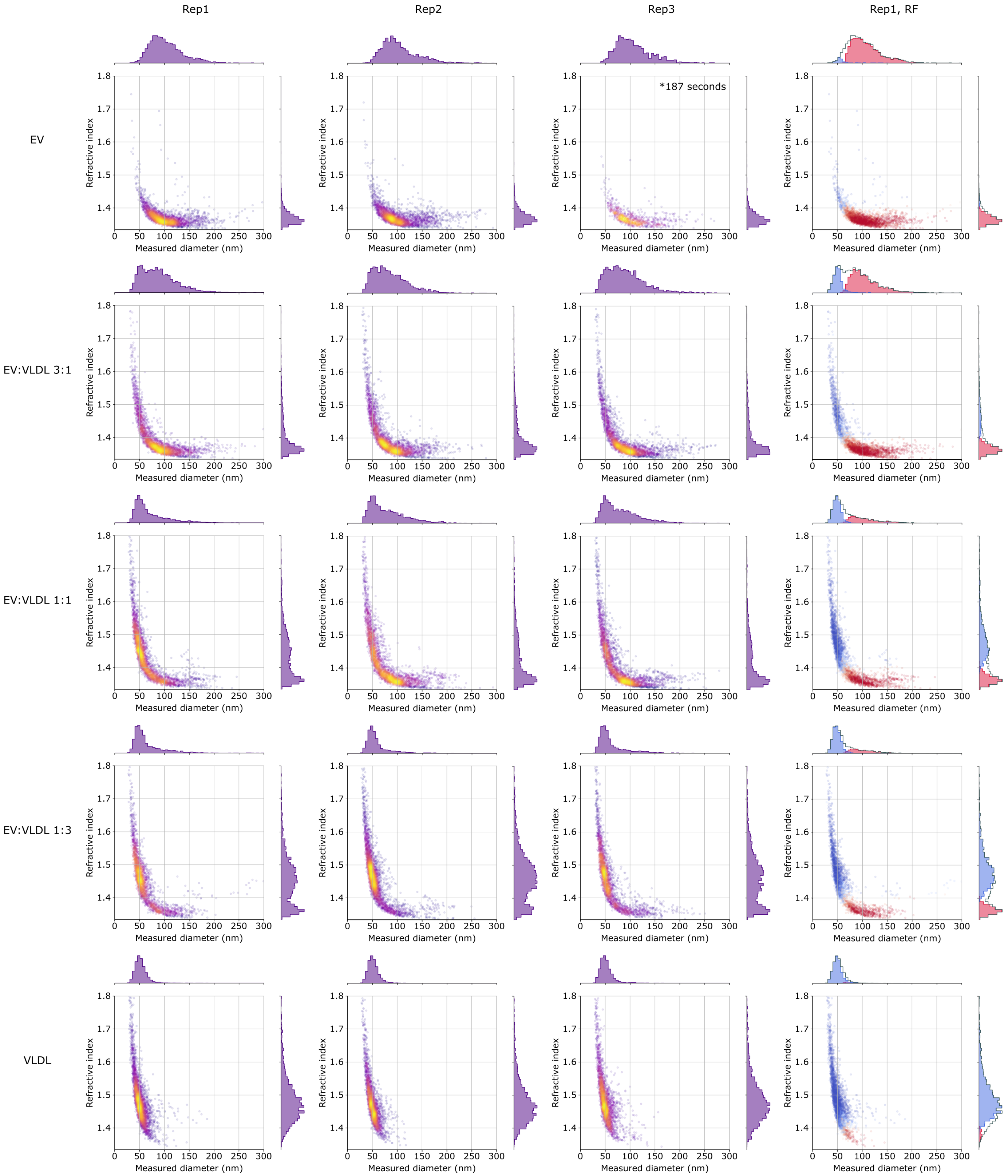


**Figure S8: iNTA measurements of EV-VLDL mixtures.** The three left columns show technical replicates of iNTA measurements of EV-VLDL mixes. All samples except for Rep3-EV were measured for 10 minutes. The right column demonstrates the application of random forest classifier to Rep 1.

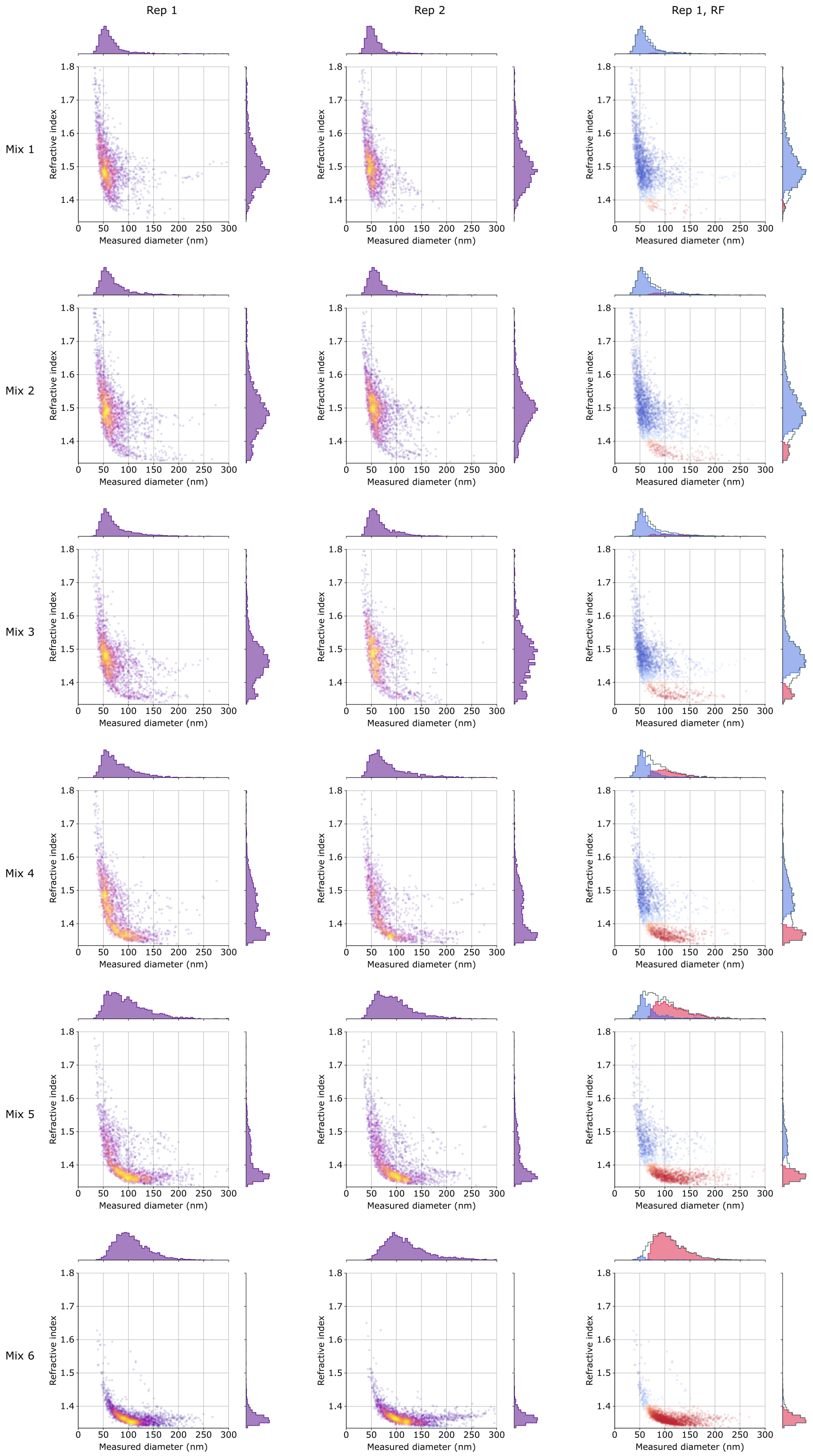


**Figure S9: iNTA measurements of EV-lipoprotein mixtures.** The two left columns show technical replicates of iNTA measurements of EV-LP mixes. All samples were measured for 10 minutes. The right column demonstrates the application of random forest classifier to Rep 1.

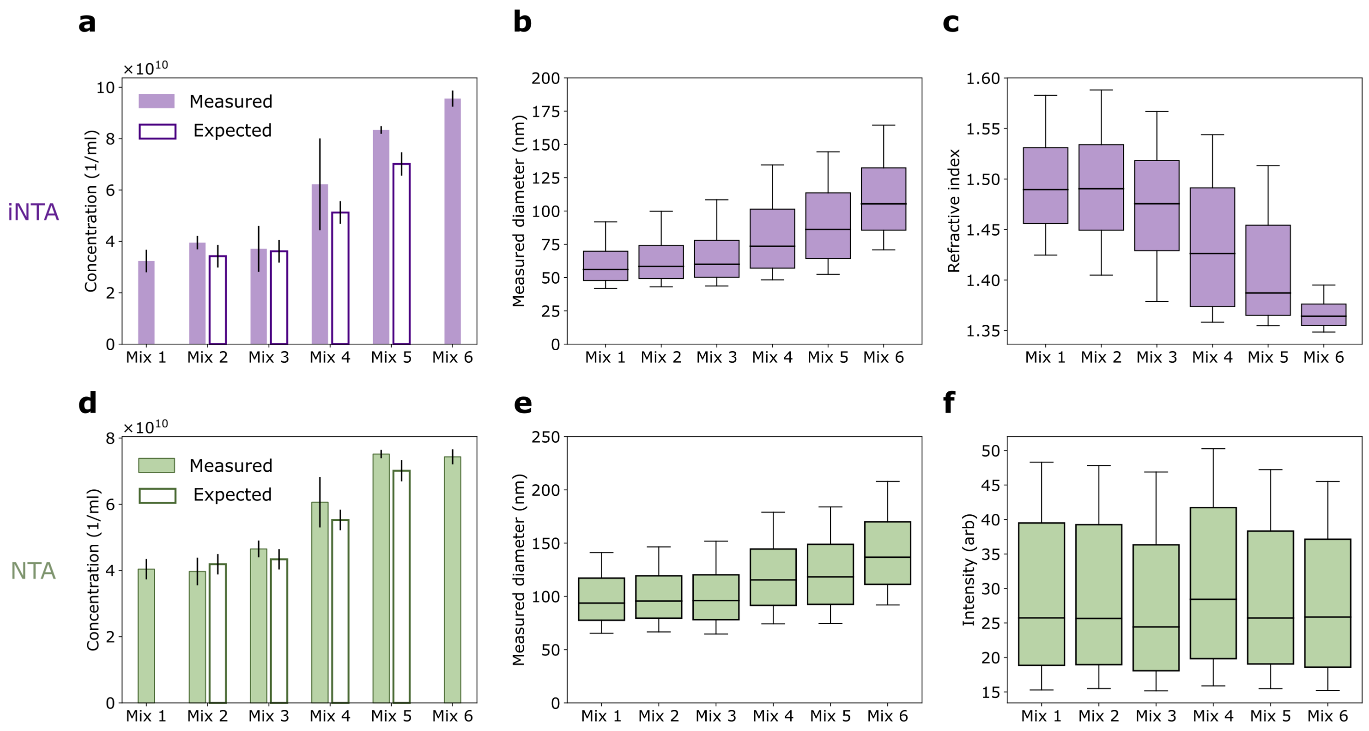


**Figure S10: Summary of iNTA and NTA measurements of EV-lipoprotein mixtures.** **(a)** Total particle concentration (mean and standard deviation), **(b)** particle diameter (median and 10-90 percentile) and **(c)** effective refractive index (median and 10-90 percentile) as measured by iNTA. **(d)** Total particle concentration (mean and standard deviation), **(e)** particle diameter (median and 10-90 percentile) and **(f)** scattering intensity (median and 10-90 percentile) as measured by NTA. The expected values in (a) and (d) are calculated using the measured values of Mix 1 (pure lipoproteins) and Mix 6 (pure EVs).

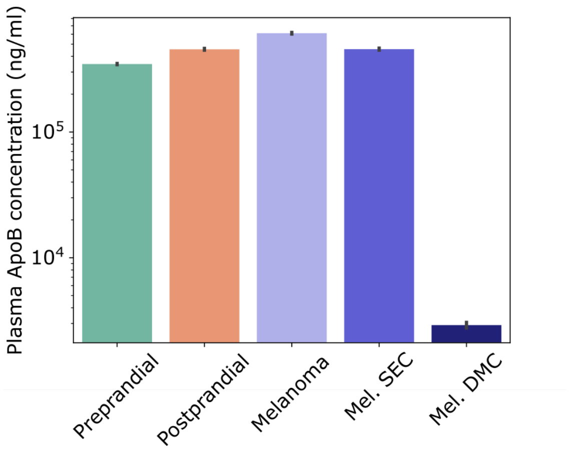


**Figure S11: ApoB content as measured by ELISA in pre- or postprandial plasma of a healthy volunteer, and lipemic melanoma patient plasma.** Mean and standard deviation of duplicate measurements are shown. Measurement of ApoB in DG processed sample was below the limit of detection.

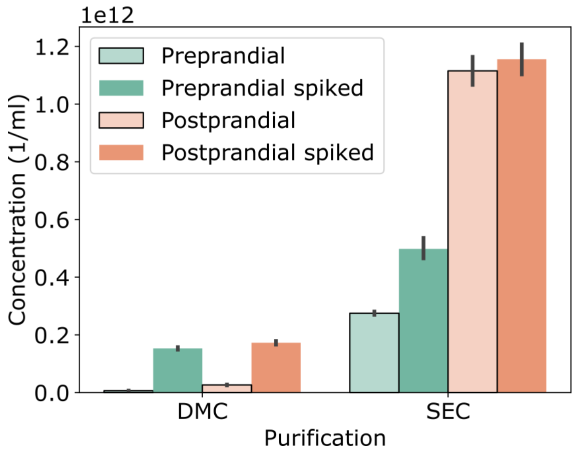


**Figure S12: NTA concentration measurements of pre- and post-prandial plasma processed by DMC or SEC, without or with addition of an exogenous EV spike.** Mean and standard deviation are shown.

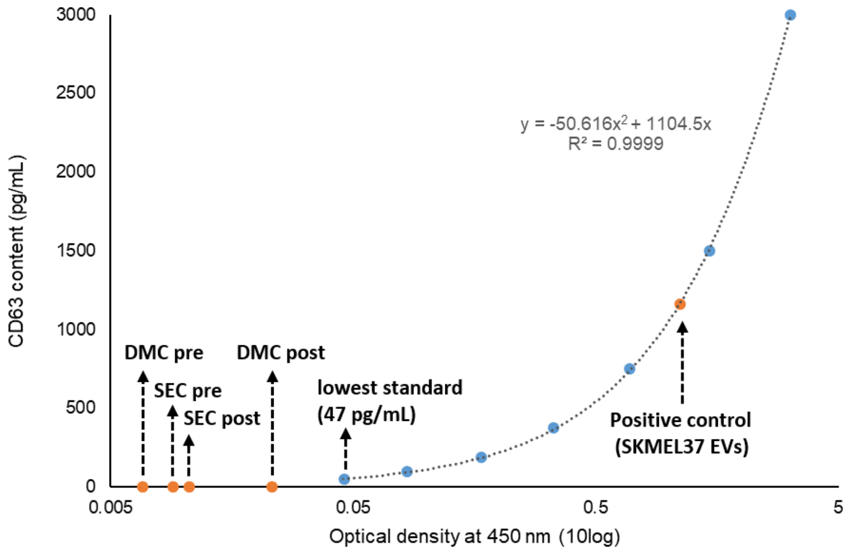


**Figure S13: CD63 ELISA.** Pre- and post-prandial plasma of a healthy volunteer was processed by SEC and DMC, and CD63 content measured as a proxy of EV presence. All samples were below limit of detection (i.e. an OD450 below the lowest standard). SKMEL-37 EVs were used as positive control.

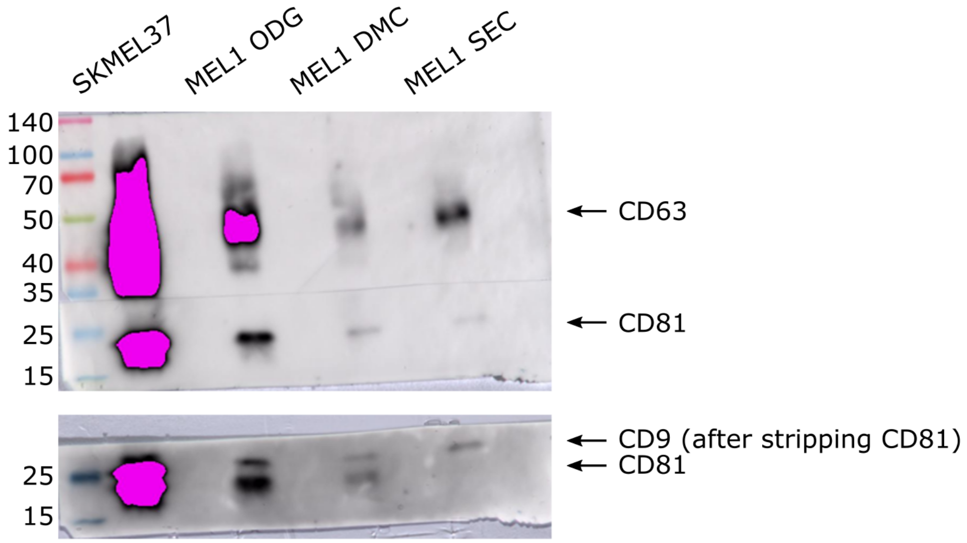


**Figure S14: Full-length Western blot corresponding to Figure 4a.**
